# Phylogenetic inference using Generative Adversarial Networks

**DOI:** 10.1101/2022.12.09.519505

**Authors:** Megan L. Smith, Matthew W. Hahn

## Abstract

**Motivation:** The application of machine learning approaches in phylogenetics has been impeded by the vast model space associated with inference. Supervised machine learning approaches require data from across this space to train models. Because of this, previous approaches have typically been limited to inferring relationships among unrooted quartets of taxa, where there are only three possible topologies. Here, we explore the potential of generative adversarial networks (GANs) to address this limitation. GANs consist of a generator and a discriminator: at each step, the generator aims to create data that is similar to real data, while the discriminator attempts to distinguish generated and real data. By using an evolutionary model as the generator, we use GANs to make evolutionary inferences. Since a new model can be considered at each iteration, heuristic searches of complex model spaces are possible. Thus, GANs offer a potential solution to the challenges of applying machine learning in phylogenetics.

**Results:** We developed phyloGAN, a GAN that infers phylogenetic relationships among species. phyloGAN takes as input a concatenated alignment, or a set of gene alignments, and infers a phylogenetic tree either considering or ignoring gene tree heterogeneity. We explored the performance of phyloGAN for up to fifteen taxa in the concatenation case and six taxa when considering gene tree heterogeneity. Error rates are relatively low in these simple cases. However, run times are slow and performance metrics suggest issues during training. Future work should explore novel architectures that may result in more stable and efficient GANs for phylogenetics.

**Availability:** phyloGAN is available on github: https://github.com/meganlsmith/phyloGAN/.

## 1 Introduction

Inferring phylogenetic relationships among species is a central goal of many studies in evolutionary biology. Advances in sequencing technologies and in methods for inferring phylogenies from these data have proliferated in recent years (Scornavacca et al., 2020). However, phylogenetic inference is complicated by several factors, including gene tree heterogeneity due to incomplete lineage sorting (ILS), introgression, horizontal gene transfer, and gene duplication and loss (Maddison, 1997), as well as substitution rate variation across sites and taxa. At present, most studies rely on concatenation approaches (which ignore gene tree heterogeneity) and/or summary approaches (which take inferred gene trees as input to infer a species tree). While both approaches have proven powerful, they suffer from distinct limitations. Concatenation approaches combine all gene sequences into a single alignment for inference and can be misled when gene tree heterogeneity is high (Kubatko and Degnan, 2007). On the other hand, summary approaches assume that gene trees have been inferred accurately, and that recombination occurs between but not within the genes used for inference (Bryant and Hahn, 2020). An ideal approach would take as input a set of sequence alignments and infer a species tree directly while considering processes affecting gene tree topologies, branch lengths, and sequence evolution. While such approaches exist (e.g.,*BEAST; Heled and Drummond, 2010), they are computationally demanding and therefore not applicable to large genomic datasets.

Machine learning has proven to be a powerful approach in population genetics when large datasets need to be analyzed under complex models (Schrider and Kern, 2018). In supervised machine learning, a model is trained using data with known labels. Since labeled training data are generally not available in population genetics, simulations are used to construct these training datasets. In a typical supervised machine learning approach to classification, simulations are performed under each model of interest, a classifier is trained to discriminate amongst these models using the simulated data, and then the classifier is applied to empirical data to select the best model. While this approach is powerful when a small number of models are considered, it requires that a large number of simulations be conducted under each model of interest prior to training the classifier. This quickly becomes computationally infeasible as model spaces become larger.

In phylogenetics, the model space is, at a minimum, the number of possible tree topologies. While there are only three possible unrooted trees for four taxa, this number sharply increases with the number of taxa: with only ten taxa there are 2,027,025 possible topologies. Because of this, supervised machine learning approaches have largely been limited to inferring relationships among quartets of taxa (e.g., Suvorov et al., 2020; Zou et al., 2020; Solís-Lemus et al., 2022). While these approaches performed well on quartets, this limited taxon sampling prevents these methods from competing with traditional approaches (Zaharias et al., 2022). Although other approaches have been proposed to apply machine learning in phylogenetics (e.g., Nesterenko et al., 2022), no current approach can infer a species tree for more than four taxa directly from DNA sequence data using a supervised machine learning framework.

The problem of a large model space in phylogenetics is not new: likelihood, parsimony, and Bayesian approaches have all confronted this challenge by using heuristic search strategies to find the optimal tree topology without visiting all possible topologies. In a heuristic search, a new tree is proposed each iteration, and, if that tree is better than the current tree based on some optimality criterion (e.g., has a higher likelihood), then the tree is accepted. One obvious path forward to applying deep learning approaches in phylogenetics is to employ similar heuristic search strategies. One candidate for this is generative adversarial networks (GANs; Good-fellow et al, 2014).

GANs have previously been applied in population genetics to simulate artificial genomes and alignments (Yelmen et al., 2021; Booker et al., 2022) and to estimate population genetic parameters (Wang et al., 2021). GANs consist of two networks: a generator and a discriminator. The goal of the generator is to generate data that could have reasonably been drawn from the same distribution as observed data, while the goal of the discriminator is to distinguish real from generated data. At the end of the training process, the generator should produce realistic datasets. Recently, a GAN was used to estimate parameters of a population genetic model (Wang et al., 2021). The generator simulated data under some parameterized population genetic model, and at each iteration, new parameter values were proposed via simulated annealing. When the GAN produced realistic data and training was halted, the current parameters of the generator were inferred to be the best parameters to explain the real data. Using a similar approach, GANs could be used to heuristically search tree space.

Here, we introduce a new model, phyloGAN, and critically evaluate its performance. phyloGAN heuristically explores phylogenetic tree space to find a tree topology that produces generated data that are similar to observed data. We introduce two versions of phyloGAN: a concatenated version and a version that accommodates gene tree heterogeneity (such as is produced by ILS). We demonstrate the use of phyloGAN on several simulated datasets and apply it to infer relationships among seven species of fungi. We evaluate the performance of phyloGAN based on error rates and several metrics useful for assessing GAN training. While phyloGAN is able to accurately infer relationships among up to ten taxa in the concatenation case and six taxa while considering gene tree heterogeneity, performance metrics indicate issues in training, and run times are slow compared to conventional approaches.

## 2 Methods

Given a tree topology and distribution of branch lengths, phyloGAN works by generating sequence data, comparing these data to real data using a convolutional neural network (CNN), and proposing new tree topologies to find a topology that produces data that are similar to the real data (Figure 1; Algorithm 1, Supporting Information). We begin with a pre-GAN phase to infer a scale parameter for an exponential distribution on branch lengths that produces generated data with the number of variable sites similar to that in the real data. Then, we can start to train phyloGAN. The generator generates tree topologies that are used as input in an evolutionary simulator, and the discriminator is a CNN that learns to distinguish generated from real data. As in Wang et al. (2021), our approach differs from a traditional GAN, in which both the generator and the discriminator are neural networks.

**Figure 1.**
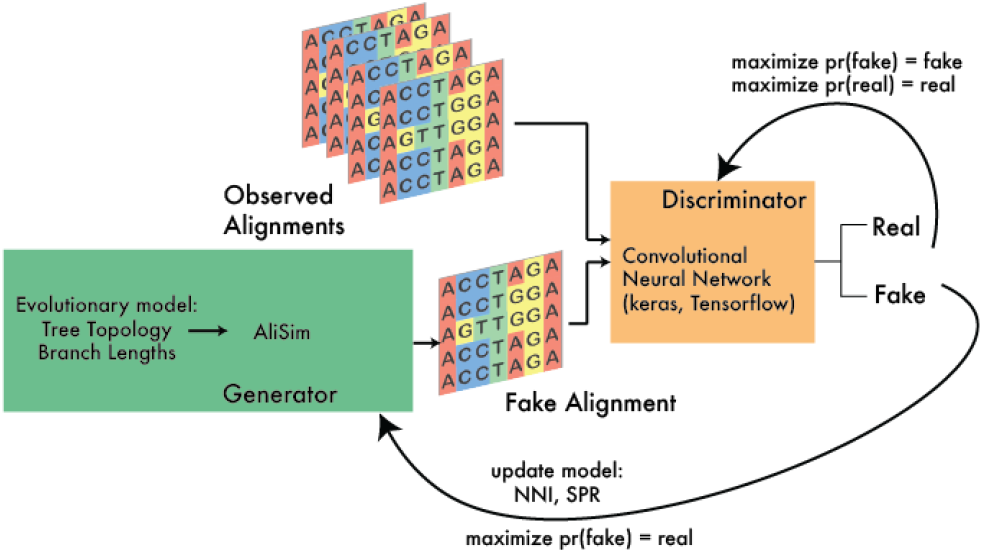
Overview of phyloGAN. The generator generates a tree topology and branch lengths, which are used as input into an evolutionary simulator (AliSim). At each iteration, new topologies are proposed using nearest neighbor interchange (NNI) and subtree pruning and regrafting (SPR). The discriminator is a CNN trained to differentiate real and generated data.

Here, we first describe the general strategy employed in phyloGAN. We then describe our pre-GAN phase to infer a reasonable parameterization on the distribution of branch lengths. Following this, we describe the generator and discriminator in more detail. Finally, we describe an extension of phyloGAN to accommodate gene tree heterogeneity.

The main input to phyloGAN is a multiple sequence alignment. Additionally, the user must provide a model of sequence evolution and bounds on the scale parameter for the exponential distribution on branch lengths. The user must also supply several arguments related to training (described in the online documentation at https://github.com/meganlsmith/phylo-GAN/). The main outputs of phyloGAN are the tree topology, τ, at each iteration, the generator and discriminator loss at each iteration, and the trained discriminator.

Training a GAN is a minimax game in which the generator aims to maximize the probability that generated data is classified as real data, and the discriminator aims to minimize the probability that either generated or real data is misclassified. Here, our generator aims to minimize the generator loss function. As in Wang et al. (2021), with *M* blocks of simulated data {*z*^*(l)*^ …*z*^*(m)*^} generated under τ, the generator loss is calculated using the cross-entropy function:

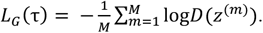

Where *D(x)* is the predicted probability that the block *x* is real. Note that, in GANs, the generator loss is calculated using the discriminator. The discriminator aims to classify generated regions as fake and real regions as real by minimizing the discriminator loss function for *M* regions of simulated data {*z*^*(l)*^,…, *z*^*(m)*^} and M regions of real data X = {*x*^*(l)*^,…, *x*^*(m)*^}. Again, as in Wang et al. (2021), the discriminator loss function is calculated using the binary cross-entropy function:

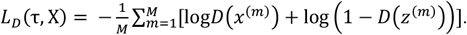

### 2.1 Stage 1: Inferring λ

During training, branch lengths are drawn from an exponential distribution with scale parameter 1/ λ. To ensure that simulated datasets are reasonably similar to the real data in terms of the number of variable sites, we precede our training by inferring a λ that produces datasets similar to the real data. We begin with some value of λ drawn from a uniform prior with bounds provided by the user. Then, at each training iteration we propose *k* new values of λ using simulated annealing. For each proposal, we simulate *M* regions of sequence data of length *L* under a random tree topology with branch lengths drawn from an exponential distribution with scale parameter 1/λ. We calculate the average proportion of invariable sites (*pinv*) across the *M* regions, and then calculate the distance between this value and the same value calculated from *M* regions of real data:

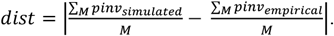

At each iteration, we keep the proposal with the smallest distance. Then, this distance is compared to the current distance. We accept the proposed value if it decreases the distance, or with some probability proportional to the ratio of the distances and the temperature, *T*. Here, *T* determines how often we accept bad proposals. *T* starts at 1 and decreases to 0 over a fixed number of iterations. Specifically, *T* is calculated as:

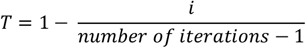

where *i* is the current iteration, indexed at zero. The inferred value of λ is used to parameterize the distribution of branch lengths during the training stage of phyloGAN.

### 2.2 The Generator

phyloGAN needs to generate tree topologies, τ, and branch lengths that can then be used as input into an evolutionary simulator. Here, we use nearest neighbor interchange (NNI) and subtree pruning and regrafting (SPR) moves to propose tree topologies. These moves are implemented using the python packages ETE3 (Huerta-Cepas et al., 2016) and Bio.Phylo (Talevich et al., 2012). Following these moves, AliSim (Ly-Trong et al., 2022) is used to generate sequence data under the proposed topologies. The proposed moves are determined by the temperature, *T*. As described above, *T* starts at 1 and decreases to 0 over a fixed number of iterations. When *T* is near 1, SPR moves are more likely, and as *T* approaches 0, NNI moves become more likely. Specifically, we draw a probability from a uniform distribution and multiply the probability by *T*. If the probability is greater than 0.4, we propose tree topologies using SPR; otherwise, we propose tree topologies using NNI. If the first proposal type (NNI or SPR) fails to produce the maximum number of trees to be considered at the step, then we use trees from the other proposal type to generate additional trees up to the desired number of trees.

Training begins either with a random start tree or with a neighbor-joining tree inferred from the real data. At each iteration, *k* new topologies are proposed using NNI and/or SPR. The generator loss is calculated for each proposed topology (with branch lengths drawn from an exponential distribution separately for each region), and the proposed topology with the lowest generator loss is retained. At the end of each iteration, the best proposed topology, τ, is always accepted if the generator loss is less than or equal to the previous iteration. Otherwise, we use the temperature *T* (calculated as described above) to determine whether to accept the proposed topology. The acceptance probability is calculated as follows:

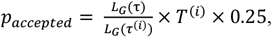

where τ ^(i)^ is the proposed topology, τ is the current topology, and *T*^*(i)*^ is the current temperature. If the proposed topology is accepted, the discriminator is trained as described below, and the new topology is used to generate the next generation of trees through NNI and SPR proposals. If the proposed topology is not accepted, then we keep the current topology τ for the next iteration, and propose the next generation of trees from this current topology.

### 2.3 The Discriminator

As the discriminator, we use a CNN implemented in keras (Chollet et al., 2015) and tensorflow (Abadi et al., 2015), with an architecture inspired by Wang et al. (2021) and Suvorov et al. (2020). Sequence data are converted to 2D NumPy arrays (Harris et al., 2020) by coding {A,T,C,G} as {0,1,2,3}. We determine the maximum number of variable sites (*S*) to include when training by sampling *M* regions of length *L* from the real data and calculating the average number of variable sites. Any simulated or real regions with more than *S* variable sites are truncated to *S* variable sites, and any regions with fewer than *S* variable sites are padded with the missing data value “4.” To ensure that all possible combinations of individuals are considered by the convolutions when constructing these arrays for training the discriminator, we sample all quartets of individuals and provide sets of rows for each quartet. For example, if there are five species in the dataset, the initial alignment consists of five rows. However, the 2D array consists of 20 rows, corresponding to all combinations of four taxa. Training datasets are created from the real dataset in a similar fashion; *M* regions of length *L* are sampled and converted to 2D NumPy arrays as above.

The discriminator consists of several layers. First, we use a convolutional layer with 10 filters with kernel size (4,1) and a stride size (2,1) with ReLU activation. Next, we apply a MaxPooling layer with pool size (1,2) and stride size (1,2). This is followed by a second convolutional layer with 10 filters, kernel size (2,1), and stride size (2,1), which is followed by another MaxPooling layer identical to the first. Then, we apply a flattening layer, a dense layer with 128 neurons, a dropout layer with a dropout rate of 0.2, another dense layer with 128 neurons, another dropout layer, and, finally, a dense layer with a single layer and linear activation, which predicts class probabilities. When training the discriminator, we use a user-specified number of epochs. For each epoch, we simulate *M* regions under the current tree topology (with branch lengths drawn from an exponential distribution separately for each region), and then train the discriminator using a gradient descent approach.

Following Wang et al. (2021) we precede training with a pre-training phase, so that the discriminator has an overall sense of the data. For a user-specified number of pre-training iterations, we generate a random tree topology under a pure birth model with branch lengths drawn from an exponential distribution with scale parameter 1/ λ. We then generate *M* regions under the tree topology and branch lengths, draw *M* regions from the real data, and pre-train the discriminator. In pre-training, we do not accept or reject proposals; at each iteration, a tree is generated, sequence data are simulated, and the discriminator is trained.

### 2.4 Extension to gene tree heterogeneity

To extend phyloGAN to consider gene tree heterogeneity, we made several modifications. First, in stage 1, we jointly infer the coalescent time and the scaling parameter, such that we are explicitly modeling gene tree heterogeneity due to ILS. Specifically, branch lengths are now parameterized as node ages drawn from an exponential distribution with the scale parameter equal to the coalescence time, and this scale parameter is inferred in a pre-GAN phase. Additionally, there is a scaling parameter associated with simulating sequence data along these gene trees that is jointly inferred in the pre-GAN phase. This parameter (the branch-scale parameter in Ali-Sim) scales the gene tree prior to simulation of sequence data and impacts the number of substitutions generated along edges of the gene tree. Finally, in order to incorporate ILS, we added a step to the simulator. For each simulated region, *m*, gene trees, *G*, are simulated within the species tree under the multispecies coalescent model using DendroPy (Sukumaran and Holder, 2010); sequence data (with alignment lengths based on the empirical data) are simulated along these gene trees in Ali-Sim, as above to generate a series of matrices for each gene tree. These matrices are concatenated into a single matrix before being used as input into the discriminator. We refer to this version as phyloGAN-ILS hereafter.

### 2.5 Simulations

To evaluate the performance of phyloGAN, we used several simulated datasets. For each combination of tree topology, sequence length, and branch lengths, we simulated ten alignments and ran phyloGAN on each alignment. All simulations of sequence data were performed using the Jukes-Cantor model (Jukes and Cantor, 1969). First, to evaluate phyloGAN’s performance over a range of sequence and branch lengths, we focused on a five-taxon tree (Figure 2). As a baseline, we set all branch lengths to range between 0.0125 and 0.0375, and evaluated performance on datasets with sequence lengths ranging from 1,000 to 1,000,000 bp. Then, we explored branch lengths from 0.1x to 10x the original values with 5,000 bp sequences. To explore the performance of phyloGAN on larger trees, we considered trees with 5-20 taxa (Figure S1), with 50,000 bp sequences and branch lengths similar to the baseline case above.

**Figure 2:**
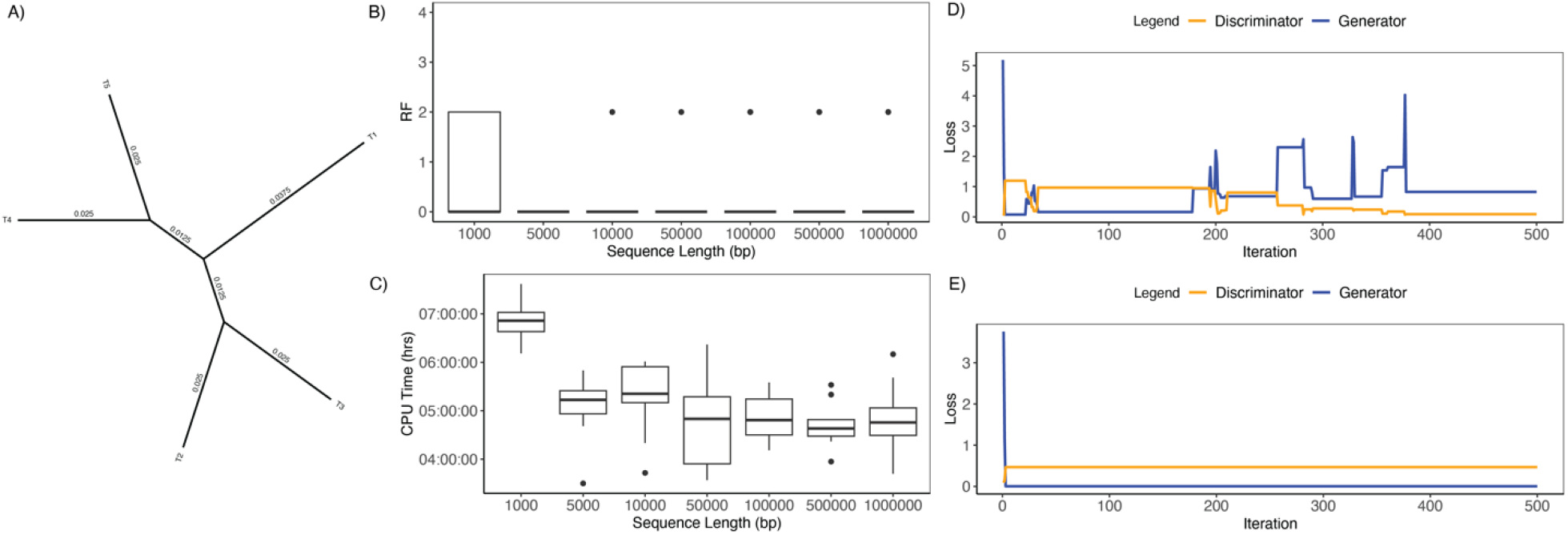
Results across sequence lengths. A) Five-taxon tree used in simulations. B) RF distance between true tree and the MCC tree. A value of 0 indicates identical trees. The maximum RF distance in this case is 4. C) CPU time (in hours) of phyloGAN. D) Generator and discriminator loss for a sequence of 10,000 bp that converged on the correct tree topology. E) Generator and discriminator loss for a sequence of 10,000 bp that failed to converge on the correct tree topology.

We ran phyloGAN on the five-taxon datasets as follows. We used a random tree as the starting tree. We considered 25 regions of 500 bp in each minibatch. We set the minimum and maximum bounds on λ to 10 and 100, respectively, for the baseline case. The bounds on λ were adjusted when branch lengths varied to ensure that the bounds encompassed the values used in simulations. Stage 1 (i.e., inferring λ) included 100 iterations and considered ten proposals per iteration. We ran 20 pre-training iterations, and trained the discriminator for 30 epochs during pre-training. We then ran phyloGAN for 500 training iterations, and trained the discriminator for 25 epochs when new topologies were accepted. At each training iteration, we considered a maximum of 10 trees. For larger trees, we increased the number of training iterations (to 1000 for six to seven taxa, 2000 for eight taxa, and 5000 for more than eight taxa). We found the maximum clade credibility (MCC) tree using the “sumtrees” command in DendroPy (Sukumaran and Holder, 2010; Sukumaran and Holder, 2015) after discarding the first 25% of trees as burn-in.

To evaluate the performance of phyloGAN-ILS, we simulated 5000 gene trees on a six-taxon species tree (Figure S2). For each gene tree, we simulated between 300 and 1200 bp of sequence data. We ran phyloGAN-ILS for 20 pretraining and 1,000 training iterations, and used a random tree as the starting tree. We trained the discriminator for 20 epochs in pretraining and 30 epochs in training, as above. For each generated region, we simulated gene trees and sequence data (with lengths based on the empirical data) until we had 1000 bp of generated data. We used a mini-batch size of 25 regions.

### 2.6 Evaluation

To evaluate the performance of phyloGAN on simulated datasets, we considered several metrics. First, we considered the Robinson-Foulds (RF) distance, which measures the distance between two trees by counting the number of splits that are unique to each of the trees (Robinson and Foulds, 1981). We measured the RF distance between the MCC tree and the true tree and between the tree at the end of a phyloGAN run (the final tree) and the true tree. We considered both generator and discriminator loss, calculated as described above. We considered discriminator accuracy on generated and training data, and the average discriminator accuracy. Discriminator accuracy was calculated after the discriminator had been trained on a set of generated and real data. Finally, we considered what we call generator accuracy. At each iteration, if a proposed tree was accepted, then, prior to training, we calculated the discriminator accuracy on the generated data. This measures the ability of the generator to produce realistic data before the discriminator has been given the opportunity to train on the generated data and thereby improve its accuracy. Ideally, generator loss should decrease throughout the run, and discriminator loss should increase slightly. Discriminator accuracy would optimally be around 0.5 on both generated and training data by the end of the run, indicating a generator that produces data highly similar to the observed data. Similarly, generator accuracy should be around 0.5 at the end of the run.

### 2.7 Empirical data

To evaluate the performance of phyloGAN on an empirical dataset, we used a subset of the fungal dataset from Rasmussen and Kellis (2011). We considered seven species: *Candida lusitaniae, C. guillermondii, Debaryomyces hansenii, C. albicans, C. tropicalis, C. parapsilosis*, and *Lodderomyces elongisporus*. We retained only those gene families that were present in exactly one copy in each sampled species, and we removed all sites with > 50% gaps in a gene alignment prior to concatenating the retained gene alignments. We ran phyloGAN as described above for simulated data, using 5000 training iterations, and found the MCC tree using DendroPy. We used a random tree as the starting tree.

## 3 Results

phyloGAN was fairly accurate in the five-taxon case (Figure 2B). For all sequence lengths greater than 1,000 bp with the baseline branch lengths, the MCC tree was identical to the true tree in at least 80% of replicates. Results were slightly worse when using the final accepted tree, rather than the MCC tree, for inference (Supporting Figure S3). Given this, we treat the MCC tree as the inferred tree below, except when noted otherwise. phyloGAN generally took longer in cases where inference was more difficult (e.g., with shorter alignments; Figure 2C). phyloGAN required an average of ∼5 CPU hours to run with 5,000 bp, and used ∼780 Mb of memory. In ∼13% of these baseline cases when more than 1,000 bp were used, phyloGAN inferred an incorrect tree topology. These cases were often easily diagnosable using metrics for assessing GAN performance. Specifically, in many of these runs the network got stuck early, and no new trees were accepted (Supporting Figure S4), so losses and accuracies do not change throughout the run (Figure 2 D,E; Supporting Figure S4). Thus, although phyloGAN sometimes fails to converge on the correct topology, these cases are often easy to diagnose.

phyloGAN was also fairly accurate across all branch lengths greater than the shortest considered here (Figure 3A), while CPU times were highest for the shortest and longest branch lengths (Figure 3B). Results were again worse when using the final accepted tree, rather than the MCC tree for inference, particularly when branch lengths were longer (Supporting Figure S5). As above, in the cases where phyloGAN did not infer the correct topology, diagnostics often indicated issues during the run. For example, in one case where the MCC tree was incorrect but the final tree was correct, accuracies, losses, and RF distances changed minimally throughout the run, suggesting issues adequately exploring tree space (Supporting Figure S6). In cases where both the MCC tree and the final tree were incorrect, metrics often indicated a failure to adequately explore tree space (Supporting Figure S7).

**Figure 3:**
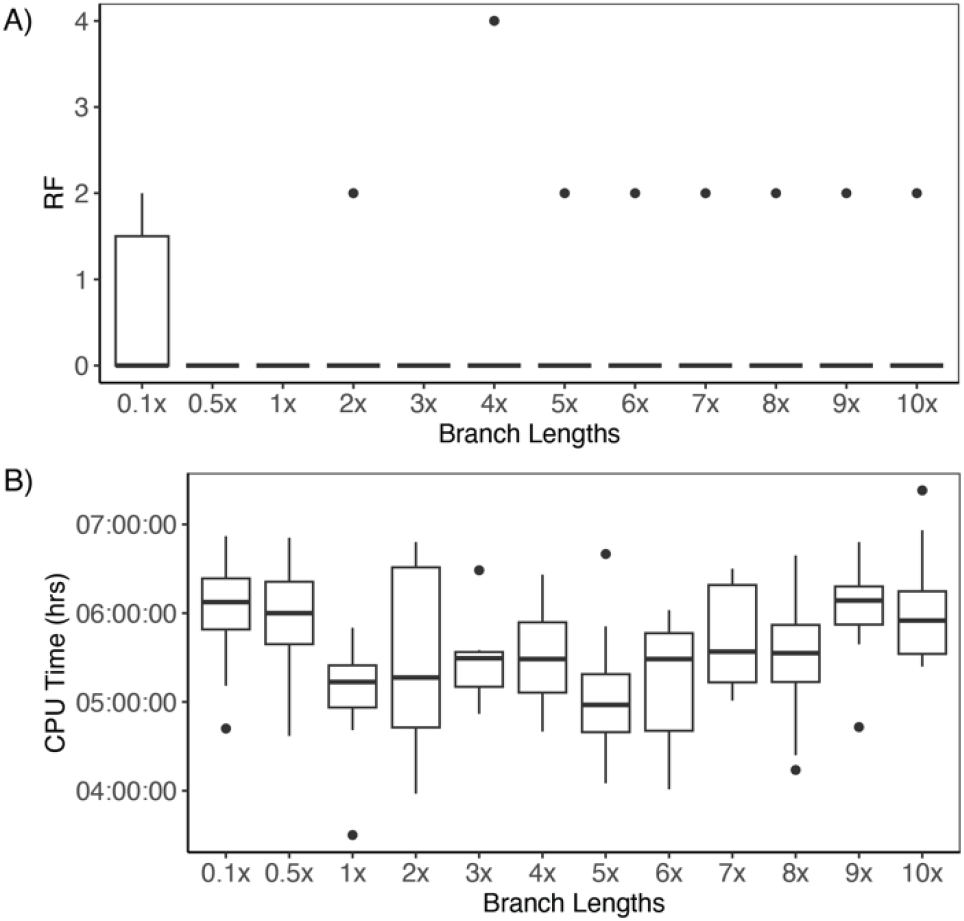
Results across branch lengths. A) RF distance between true tree and the MCC tree. A value of zero indicates identical trees. The maximum RF distance in this case is 4.B) CPU time (in hours) of phyloGAN. Branch lengths are reported in units relative to the baseline values shown in Figure 2, with 0.1x indicating branch lengths 0.1x as long as the baseline branch lengths.

phyloGAN performed well with as many as ten taxa, but accuracy began to decrease with fifteen taxa (Figure 4), and the CPU time scaled with the number of taxa. Results were again worse when relying on the final tree rather than the MCC tree for inference (Supporting Figure S8). phy-loGAN-ILS inferred the correct species tree in 100% of runs, but was substantially slower than the concatenated version (average wall time ∼ 4.6 days). phyloGAN-ILS also performed worse when relying on the final tree instead of the MCC tree, inferring the correct tree in 70% of runs.

**Figure 4.**
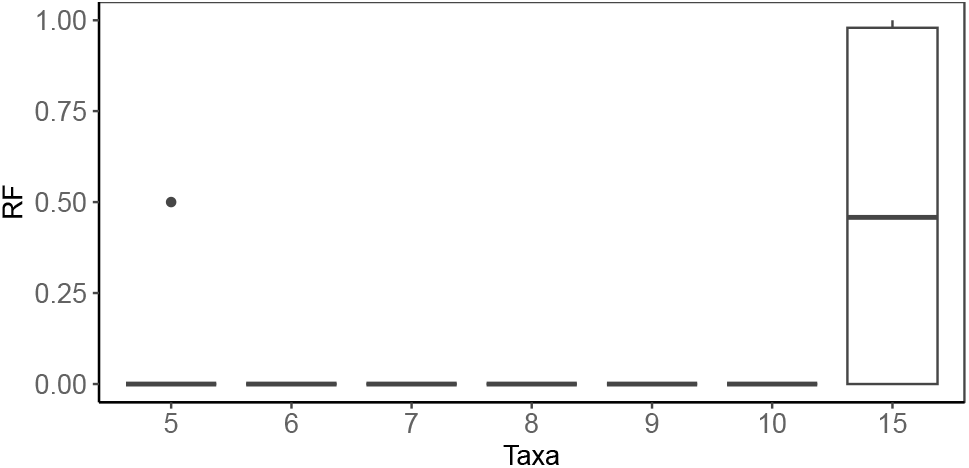
Results on larger trees. Normalized RF distances between the true tree and the MCC tree. The normalized RF distance is the RF distance normalized by the maximum RF distance for trees of the specified size. It is bounded between zero (identical trees) and one.

The MCC tree inferred using phyloGAN for the yeast species differed from the generally accepted species tree in its placement of *C. guiller-mondii* sister to *C. lusitaniae*, rather than *D. hansenii* (Figure 5). The final tree inferred using phyloGAN additionally differed from the generally accepted tree in its placement of *C. parapsiolosis* as sister to a clade including *C. tropicalis, C. albicans*, and *L. elongisporus* rather than as sister to *L. elongisporus* (Supporting Figure S13).

**Figure 5:**
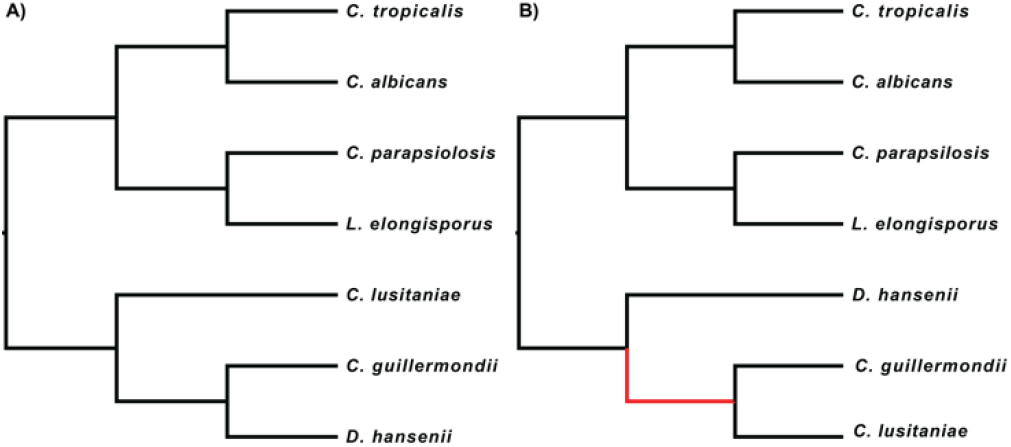
Results from fungal dataset. A) Tree topology from Rasmussen and Kellis (2011). B) MCC tree estimated in phyloGAN.

We evaluated several metrics that could be useful in diagnosing performance issues, including generator and discriminator loss and accuracy. These metrics are extremely useful in diagnosing when phyloGAN runs have failed to explore tree space (Figure 2, Supporting Figures S4, S6, S7), as discussed above. In an ideal scenario, generator loss should decrease throughout a GAN run, and discriminator loss should increase to a point. However, this is not the trend we see with phyloGAN. In successful runs, we generally see that generator loss decreases early in the run (Figure 2, Supporting Figures S6, S7, S9-11, S12). However, often the initial decrease is followed by a slow increase in generator loss, likely reflecting improvements in the discriminator. Discriminator loss quickly reaches low levels, suggesting that the discriminator learns quickly to discriminate real from generated data (Figure 2, Supporting Figures S6, S7, S9-11, S12). Ideally, discriminator accuracy should increase early in the run and then decrease to ∼0.5, since generated data are becoming more similar to real data; however, with phyloGAN, discriminator accuracy increases quickly and remains high throughout the run (Supporting Figures S4, S6, S7, S9-11, S12). Generator accuracy, however, appears to often perform as expected. It increases slowly and never reaches 1 for many replicates (Supporting Figures S4, S6, S7, S9-11, S12). Our results are more concerning for the analysis of the yeast data. Generator loss remains high, and both discriminator and generator accuracies remain high throughout the run (Supporting Figure S14). This suggests that, for this empirical dataset, our generator fails to produce realistic data.

## 4 Discussion

Using GANs to heuristically explore model spaces has previously been attempted in population genetic inference with respect to continuous parameters (Wang et al., 2021). Here, we take this approach to explore the discrete tree space associated with phylogenetic inference. We evaluated phyloGAN’s performance using simulated datasets both with and without considering incomplete lineage sorting. Error rates were low for up to ten taxa in the concatenation setting, and runs that failed to converge were often easily identified using metrics reported by phyloGAN.

phyloGAN is the first deep learning approach that can infer a phylogeny directly from sequence data for more than four taxa. While this is promising, there are several obvious limitations to our approach. First, phyloGAN’s performance begins to decrease when fifteen taxa are considered, even though this case is relatively easy using standard inference approaches. For example, we reconstructed maximum likelihood trees for the ten simulated datasets with 15 taxa using IQ-Tree (Nguyen et al., 2015) with the model of nucleotide substitution selected by ModelFinder (Kalyaanamoorthy et al., 2017) and inferred the correct tree in all replicates. Furthermore, standard approaches are much faster than phyloGAN. Even in the simplest case explored here, phyloGAN required 4-5 hours of computational time. Promisingly, phyloGAN-ILS was able to consider a more complex model space. However, this came at the cost of drastically increased run times (∼4.6 days). While this suggests that phyloGAN is not currently a competitive tool for phylogenetic inference, these results demonstrate that it is possible to design a GAN that explores phylogenetic model spaces. Future implementations may find ways to explore these spaces more efficiently.

The required computation time is perhaps the most notable limitation of phyloGAN. Computation times could be improved to an extent by adjustments to the current architecture. For example, the current architecture relies on including all combinations of quartets in training, which leads to decreased computational efficiency. Future work exploring permutation-invariant architectures (e.g., Sanchez et al., 2021) may lead to increased efficiency and allow phyloGAN to run faster with more taxa. One major limitation of GANs compared to other machine learning architectures is the requirement that they be retrained for each empirical dataset. However, although in theory other machine learning approaches allow for training a network once and applying it to multiple empirical datasets, this is challenging in practice. Most networks that have been used in phylogenetics and population genetics are not invariant to the number of samples or sites, and training data are generally simulated under an evolutionary model designed with a specific system and empirical dataset in mind. This has, in practice, severely limited the utility of networks trained on one dataset when analyzing new datasets in other machine learning approaches.

We explored several metrics for assessing GAN performance. Both loss and accuracy metrics proved useful for detecting phyloGAN runs that had failed to converge on the correct tree when analyzing simulated datasets. However, these metrics did not behave exactly as expected in an ideal scenario. Specifically, while in an ideal GAN discriminator accuracy would be near 0.5 by the end of the run, our discriminator accuracy was almost always near 1. This suggests that, even at the end of the run, the discriminator could easily distinguish real from generated data, although generator accuracy was generally below 1, suggesting some difficulties in classification of datasets that the discriminator had not yet seen. One explanation for this could be our approach to branch lengths. We do not infer branch lengths, but rather integrate over branch lengths. Therefore, the real data is always a single dataset generated under a specified set of branch lengths, while the generated data are generated under a tree topology and several sampled branch lengths. This may facilitate discrimination between the real and generated datasets, and prevent accuracy from decreasing to 0.5. While we could not easily explore the effects of inferring branch lengths, we evaluated whether metrics improved when we generated our simulated data under a distribution of branch lengths (as done for training data). While in theory this should lead to a closer match between observed and generated data, we found very similar performance to the baseline case (Supporting Figure S15). Additionally, our discriminator takes as input sequences formatted as images. Notably, no two datasets generated under the same evolutionary model (topology) would be expected to be identical—the goal is for the discriminator to recognize similar patterns in datasets generated under similar evolutionary models. While the discriminator must accomplish this goal to some extent in order for this approach to work at all, it could be more efficient to summarize the data in a way that made these similarities more obvious (e.g., using site pattern frequencies). Another potential explanation for this is that our GAN differs in a notable way from traditional GANs: in a traditional GAN, both the discriminator and the generator are neural networks. However, in phyloGAN only the discriminator is a neural network (see also Wang et al. 2021). This may lead to some differences in the performance of these metrics compared to a traditional GAN. This difference also brings into question whether these approaches are appropriately classified as GANs. However, other aspects of the architecture are identical to traditional GANs, and much of the research and terminology related to GANs remains relevant, even in the absence of a neural network as the generator. Finally, some of these limitations could be due to a suboptimal network architecture. We explored several variations during development, but found minimal improvements over the architecture presented here. For example, our kernel size does not take advantage of information across sites. To explore whether this limited phyloGAN’s performance, we considered two networks with discriminators with larger kernel sizes (one with kernel sizes (4,4) and (2,2) for convolutional layers 1 and 2, respectively and one with kernel sizes (4,2) and (2,2) for convolutional layers 1 and 2, respectively). These networks performed slightly worse than phyloGAN in a simple case (80% accuracy with 1x branch lengths and 5000 bp of data). However, there are many possible combinations of hyperparameters that could be explored, and some combinations could lead to improved performance.

Here, we focused on inferring a species tree directly from sequence data. While this approach allows us to avoid many of the limitations associated with summary approaches (i.e., approaches that infer a species tree from gene trees), inference could be improved by incorporating both sequence data and inferred gene trees. Recent studies have explored approaches for encoding phylogenetic trees as data in machine learning frameworks (Voznica et al., 2022; Sophia et al., 2022; Rosenzweig et al., 2022). Future explorations of GANs in phylogenetics may benefit from considering gene tree structures in lieu of or in addition to sequence data. While the limitations discussed here suggest limited applications of phyloGAN, our work demonstrates the potential of machine learning in phylogenetic contexts. Future work exploring alternative data structures and alternative GAN architectures (or other machine learning architectures) may pave the way to more accurate and efficient GANs for phylogenetics. Additionally, GANs are extremely flexible and can consider any process under which we can generate data. Thus, future work may explore the use of GANs while considering other processes that complicate phylogenetic inference, including both gene duplication and loss and introgression.

## Acknowledgements

We thank Daniel Schrider and members of the Schrider lab for helpful discussion. We thank Rob Lanfear, Franz Baumdicker, and two anonymous reviewers for helpful comments on the manuscript.

