## Supporting Information for "Phylogenetic inference using Generative Adversarial Networks"

| **Algorithm 1: Training phyloGAN.** | | | | | |
| --- | --- | --- | --- | --- | --- |
| Stage 1: Inferring λ | | | | | |
|  | **Input:** concatenated alignment, priors, and other arguments | | | | |
|  | **Output:** optimal λ | | | | |
| Initialize λ, a value drawn from a uniform prior distribution, and set *dist_current_*  to ∞. | | | | | |
| **for** *each training iteration i* **do** | | | | | |
|  | Set *dist_proposal_* to ∞.  **for** *each proposal k* **do** | | | | |
|  |  | Propose a new λ using simulated annealing, and generate *N* chunks of sequence data of length *L* under random tree topologies with branch lengths drawn from an exponential distribution with scale parameter $\frac{1}{\lambda}$.  **if** $\left\vert\frac{\sum_{N} p{inv}_{simulated}}{N}- \frac{\sum_{N} {pinv}_{empirical}}{N} \right\vert<$*dist_proposal_*  **do** | | | |
|  |  |  | | | Update *λ_proposal_* and *dist_current._* |
|  | **end** | | | | |
|  | **if** *dist_proposal_ < dist_current_* **do** | | | | |
|  |  | Update *λ_current_* and *dist_current_*_._ | | | |
|  | **else** with some probability (dist_current_/ dist_proposal_* Temperature) | | | | |
|  |  | Update *λ_current_* and *dist_current_*_._ | | | |
| **end** | | | | | |
| Stage 2: Inferring 𝜏 | | | | | |
|  | **Input:** concatenated alignment, priors, and other arguments | | | | |
|  | **Output:** optimal 𝜏 | | | | |
| Select the maximum number of SNPs to keep per chunk of length *L* based on the empirical alignment.  **for** *num pre-training iterations* **do** | | | | | |
|  | Generate a tree topology under a birth-death model with branch lengths drawn from an exponential distribution with the scale parameter $\frac{1}{\lambda}$(with λ set to the value inferred in Stage 1). | | | | |
|  | Train discriminator using real data and generated data. | | | | |
| **end** | | | | | |
| Initialize 𝜏 to either a random tree topology or a neighbor joining starting tree, and set *loss_current_* to ∞. | | | | | |
| **for** *each training iteration i* **do** | | | | | |
|  | Use NNI and SPR to propose *k* new tree topologies. | | | | |
|  | Set *loss_proposal_* to ∞. | | | | |
|  | **for** *each proposed topology k* **do** | | | | |
|  |  | Generate *M* chunks of length *L,* with branch lengths for each chunk drawn from an exponential distribution with scale parameter $\frac{1}{\lambda}$. | | | |
|  |  | Calculate the generator loss. | | | |
|  |  | **if** loss < *loss_proposal_* **do** | | | |
|  |  |  | | Update 𝜏_proposal_ and *loss_proposal._* | |
|  | **end** | | | | |
|  | **if** *loss_proposal_ < loss_current_* **do** | | | | |
|  |  | Accept 𝜏_current_ and *loss_current._* | | | |
|  | **else** with probability (loss_current_/ loss_proposal_ * Temperature * 0.25) | | | | |
|  |  | Accept 𝜏_current_ and *loss_current._* | | | |
|  | **if** *accept* **do** | | | | |
|  |  | **for** *num mini-batches* **do** | | | |
|  |  |  | Sample mini-batch of *M* regions X = {*x*^(1)^, …, *x*^(^*^M^*^)^}from the real data. | | |
|  |  |  | Simulate mini-batch of *M* regions {*z*^(1)^, …, *z*^(^*^M^*^)^} under 𝜏_current._ | | |
|  |  |  | Train discriminator using real data and generated data. | | |
|  |  | **end** | | | |
| **end** | | | | | |

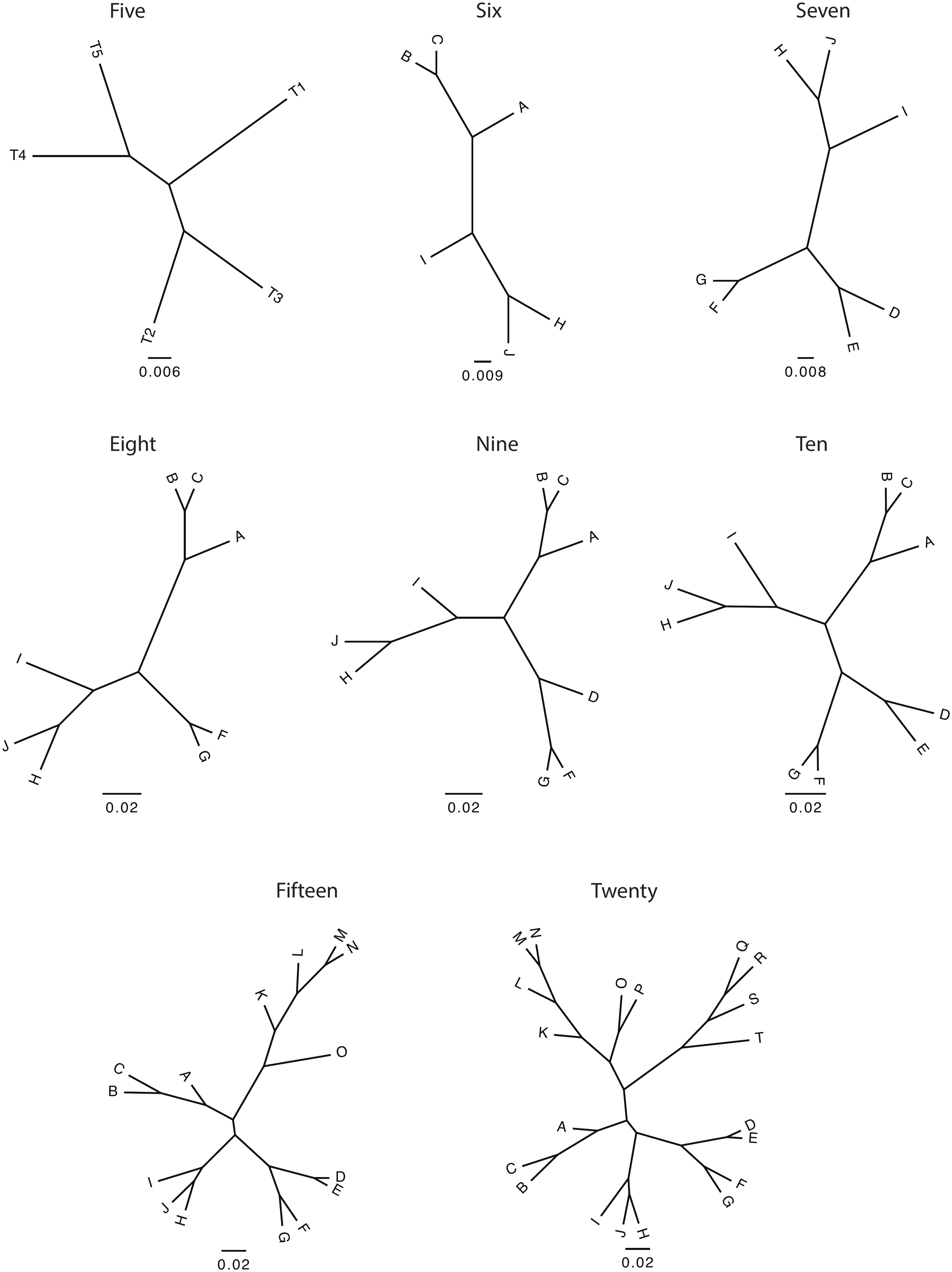

**Supporting Figure S1.** Trees used to evaluate phyloGAN performance, with number of tips shown above each.

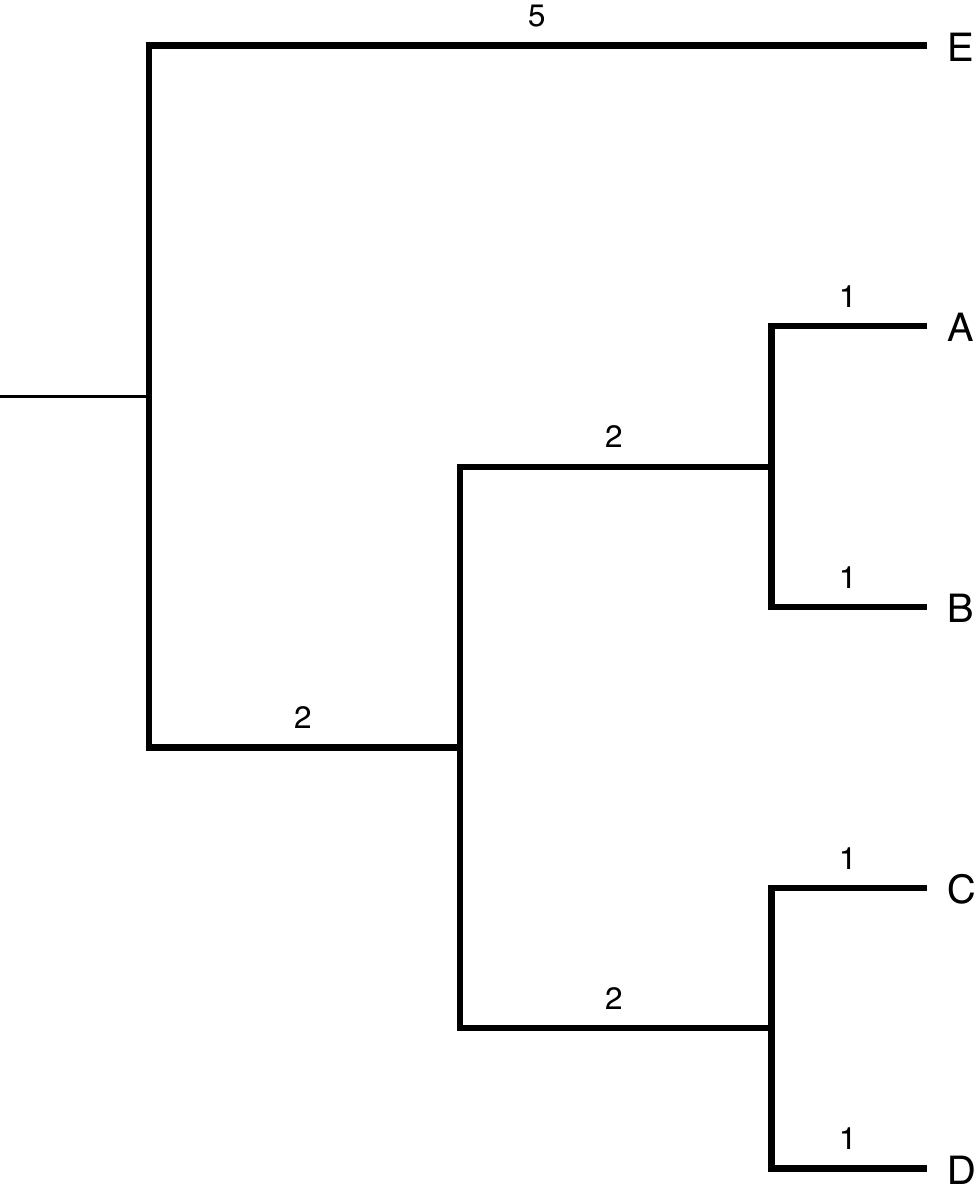

**Supporting Figure S2.** Species tree used to evaluate phyloGAN-ILS performance.

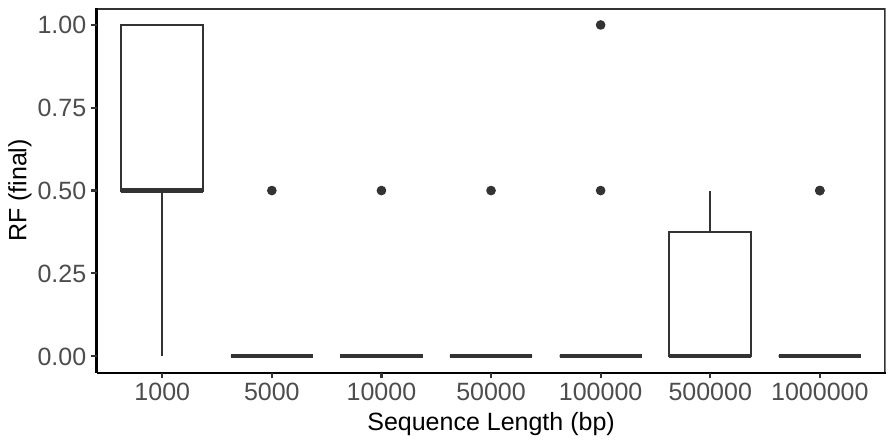

**Supporting Figure S3:** Results with baseline branch length across sequence lengths when using the final tree for inference. RF (final) is the normalized RF distance between the true and final trees.

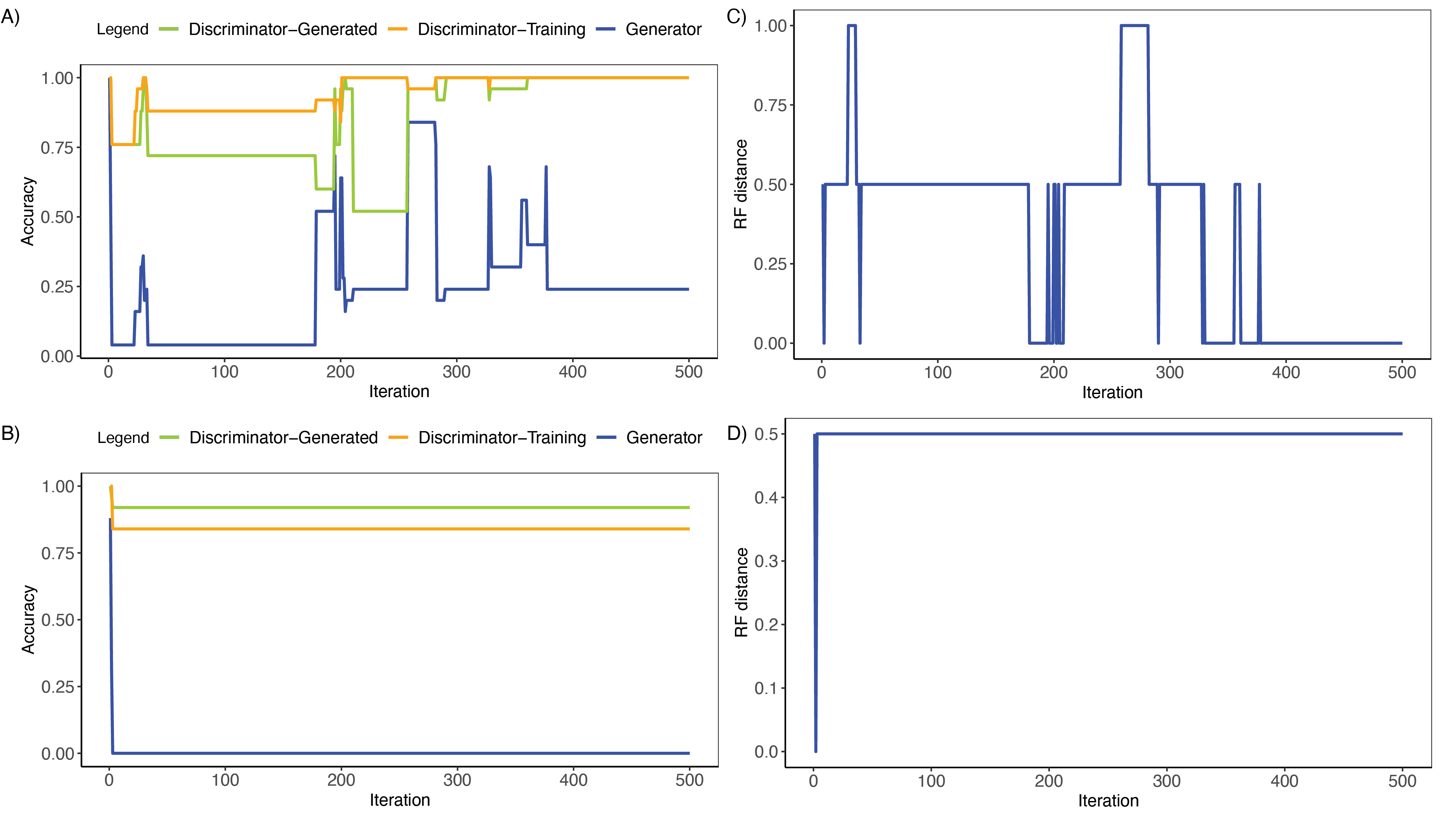

**Supporting Figure S4:** Results from baseline branch lengths and 10,000 bp. A) Generator and discriminator accuracy for a replicate that converged on the correct tree topology. Discriminator-generated and discriminator-training indicate discriminator accuracy on generated and training data, respectively, after the discriminator was trained. Generator accuracy indicates the accuracy of the discriminator on generated data prior to training for the current iteration. B) Generator and discriminator accuracy for a replicate that failed to converge on the correct tree topology. C) Normalized RF distances throughout the run for a replicate that converged on the correct tree topology. D) Normalized RF distances for a replicate that failed to converge on the correct tree topology.

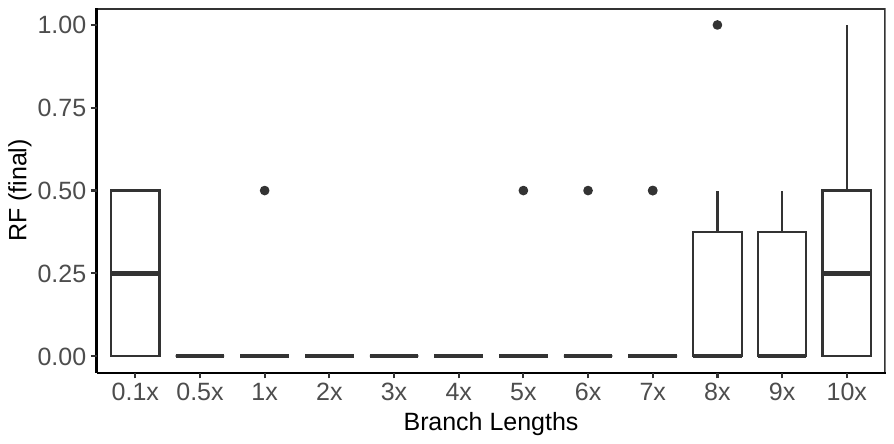

**Supporting Figure S5:** Results across branch lengths when using the final tree for inference. RF (final) is the normalized RF distance between the true and final trees.

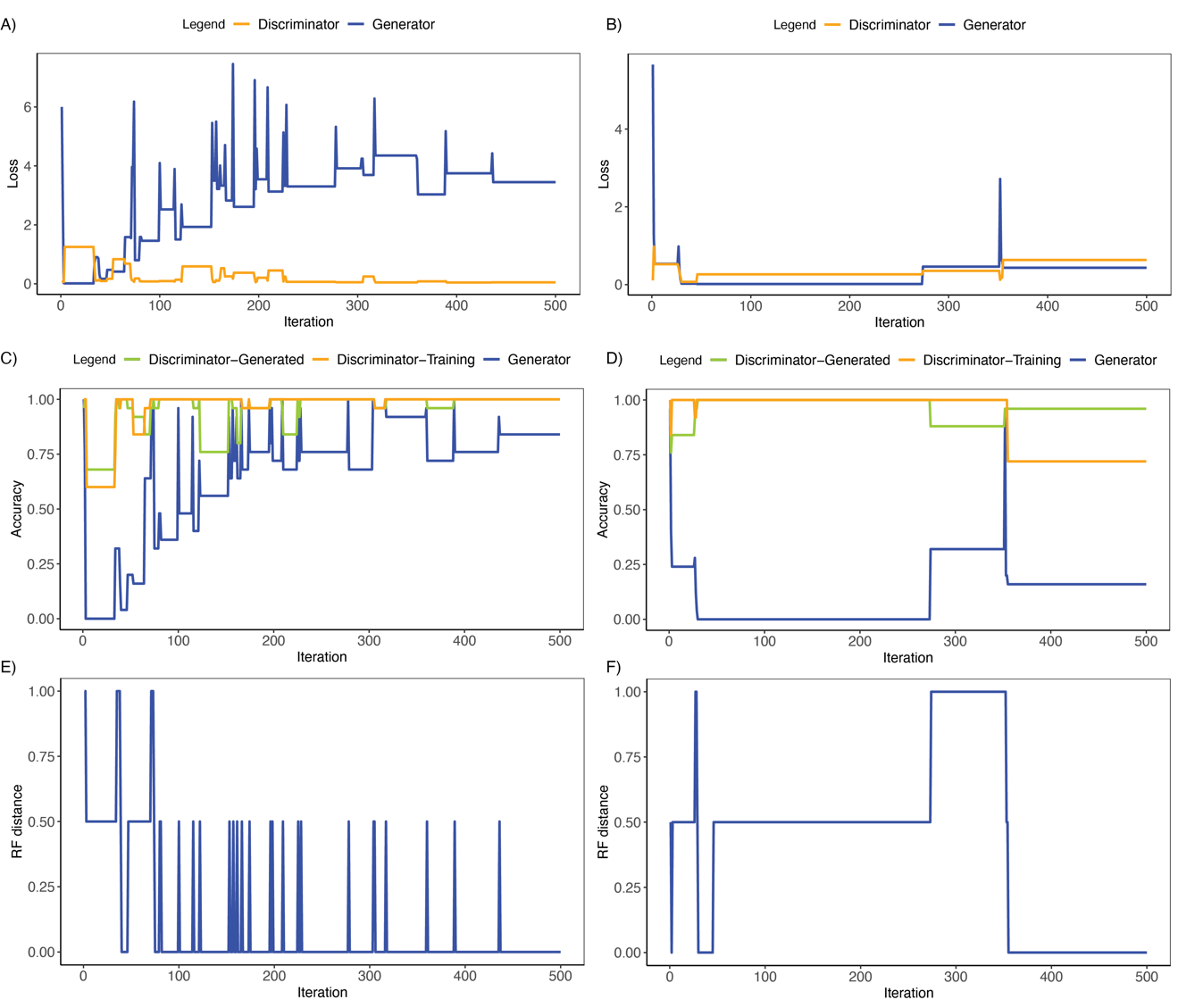

**Supporting Figure S6:** Results from 2x branch lengths and 5,000 bp from a replicate that converged on the correct topology after exploring tree space adequately and a replicate that converged on the correct topology but with less movement through tree space. A) Generator and discriminator loss for a replicate that converged on the correct tree topology. B) Generator and discriminator loss for a replicate that failed to explore tree space before converging. C) Generator and discriminator accuracy for a replicate that converged on the correct tree topology. Discriminator-generated and discriminator-training indicate discriminator accuracy on generated and training data, respectively, after the discriminator was trained. Generator accuracy indicates the accuracy of the discriminator on generated data prior to training for the current iteration. D) Generator and discriminator accuracy for a replicate that failed to explore tree space before converging. E) Normalized RF distances throughout the run for a replicate that converged on the correct tree topology. F) Normalized RF distances for a replicate that failed to explore tree space before converging.

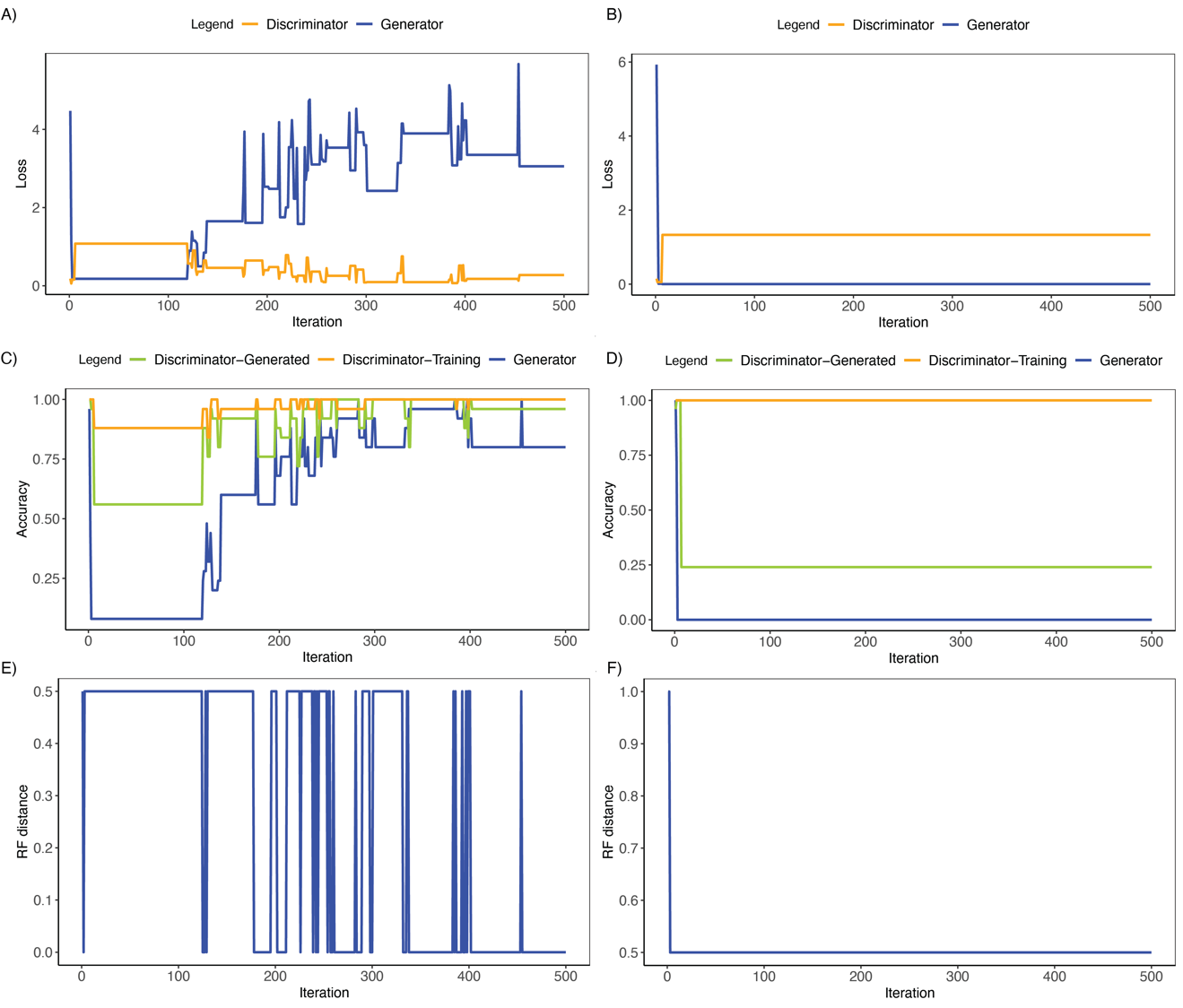

**Supporting Figure S7:** Results from 5x branch lengths and 5,000 bp from a replicate that converged on the correct topology after exploring tree space adequately and a replicate that failed to converge on the correct topology. A) Generator and discriminator loss for a replicate that converged on the correct tree topology. B) Generator and discriminator loss for a replicate that failed to converge on the correct topology. C) Generator and discriminator accuracy for a replicate that converged on the correct tree topology. Discriminator-generated and discriminator-training indicate discriminator accuracy on generated and training data, respectively, after the discriminator was trained. Generator accuracy indicates the accuracy of the discriminator on generated data prior to training for the current iteration. D) Generator and discriminator accuracy for a replicate that failed to converge on the correct topology. E) Normalized RF distances throughout the run for a replicate that converged on the correct tree topology. F) Normalized RF distances for a replicate that failed to converge on the correct topology.

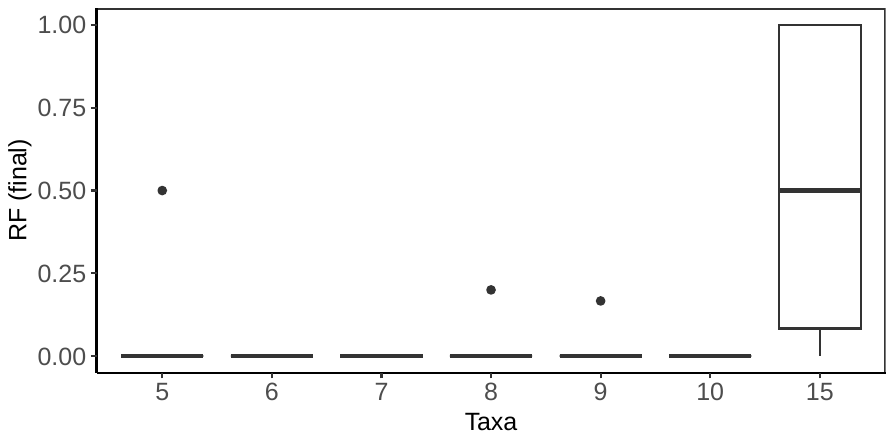

**Supporting Figure S8:** Results across tree sizes when using the final tree for inference. RF (final) is the normalized RF distance between the true and final trees.

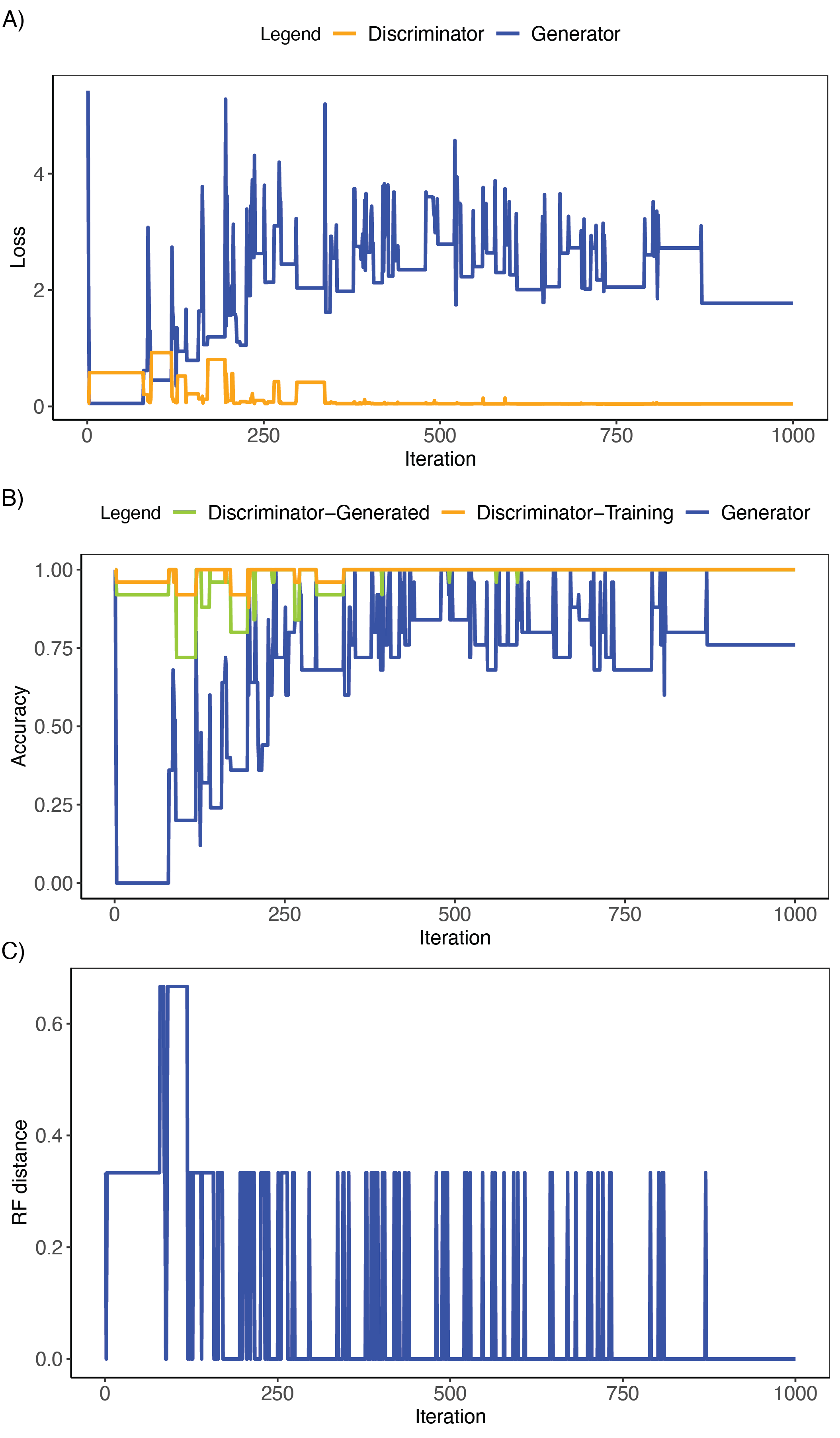

**Supporting Figure S9:** Results from phyloGAN with six taxa for a replicate that converged on the correct topology. A) Generator and discriminator loss. B) Generator and discriminator accuracy. Discriminator-generated and discriminator-training indicate discriminator accuracy on generated and training data, respectively, after the discriminator was trained. Generator accuracy indicates the accuracy of the discriminator on generated data prior to training for the current iteration. C) Normalized RF distances throughout the run.

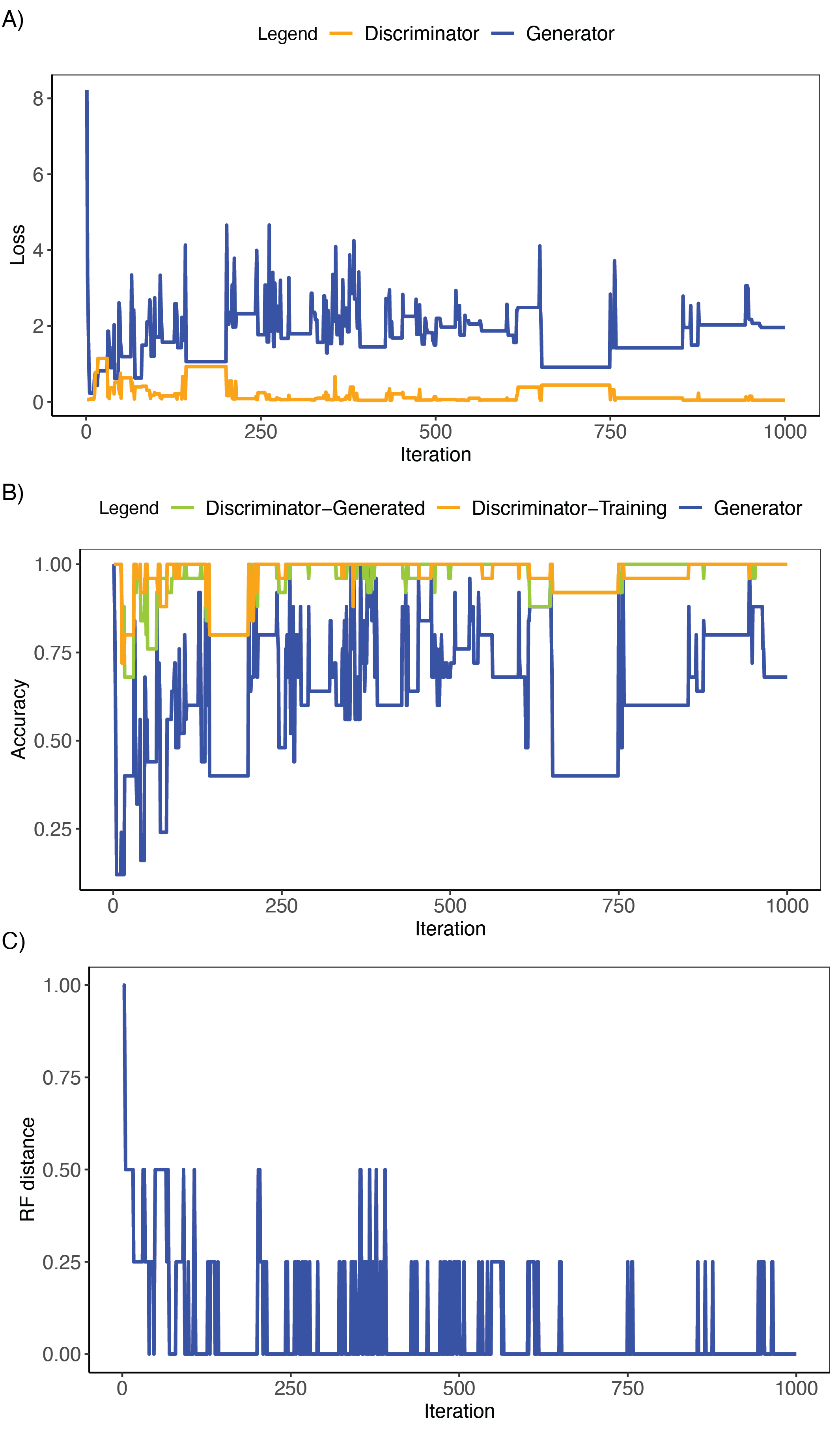

**Supporting Figure S10:** Results from phyloGAN with seven taxa for a replicate that converged on the correct topology. A) Generator and discriminator loss. B) Generator and discriminator accuracy. Discriminator-generated and discriminator-training indicate discriminator accuracy on generated and training data, respectively, after the discriminator was trained. Generator accuracy indicates the accuracy of the discriminator on generated data prior to training for the current iteration. C) Normalized RF distances throughout the run.

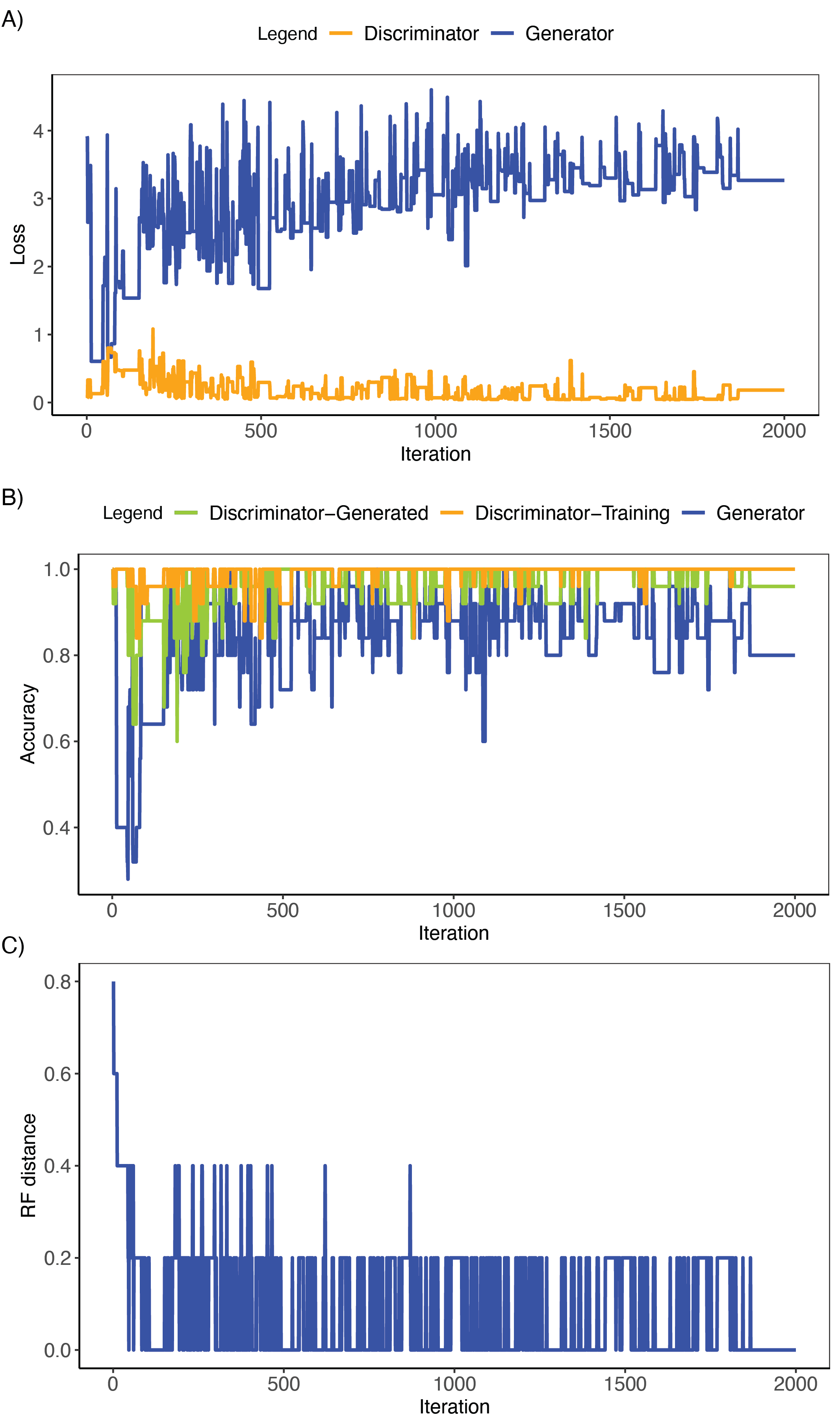

**Supporting Figure S11:** Results from phyloGAN with eight taxa for a replicate that converged on the correct topology. A) Generator and discriminator loss. B) Generator and discriminator accuracy. Discriminator-generated and discriminator-training indicate discriminator accuracy on generated and training data, respectively, after the discriminator was trained. Generator accuracy indicates the accuracy of the discriminator on generated data prior to training for the current iteration. C) Normalized RF distances throughout the run.

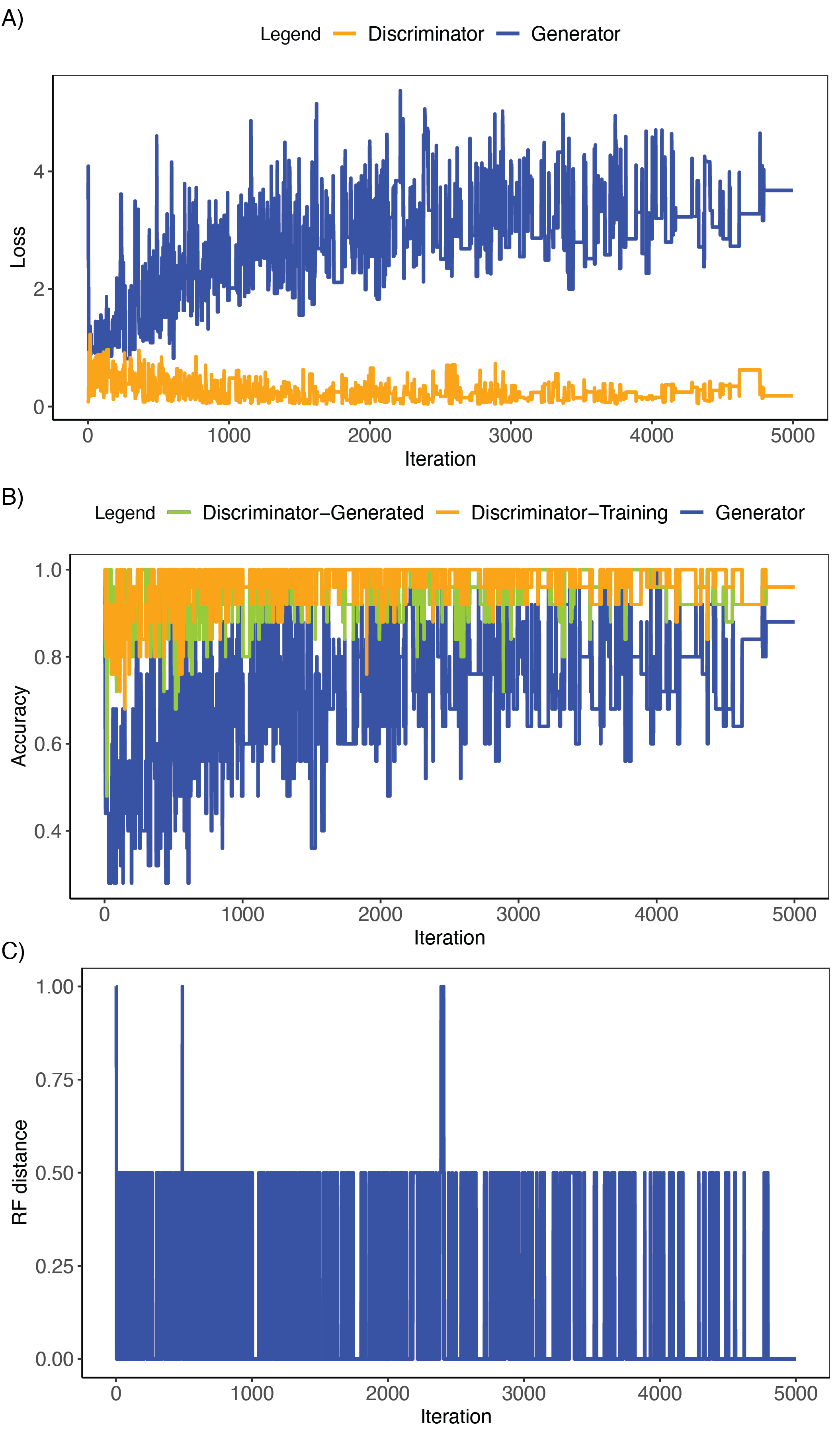

**Supporting Figure S12:** Results from phyloGAN-ILS for a replicate that converged on the correct topology. A) Generator and discriminator loss. B) Generator and discriminator accuracy. Discriminator-generated and discriminator-training indicate discriminator accuracy on generated and training data, respectively, after the discriminator was trained. Generator accuracy indicates the accuracy of the discriminator on generated data prior to training for the current iteration. C) Normalized RF distances throughout the run.

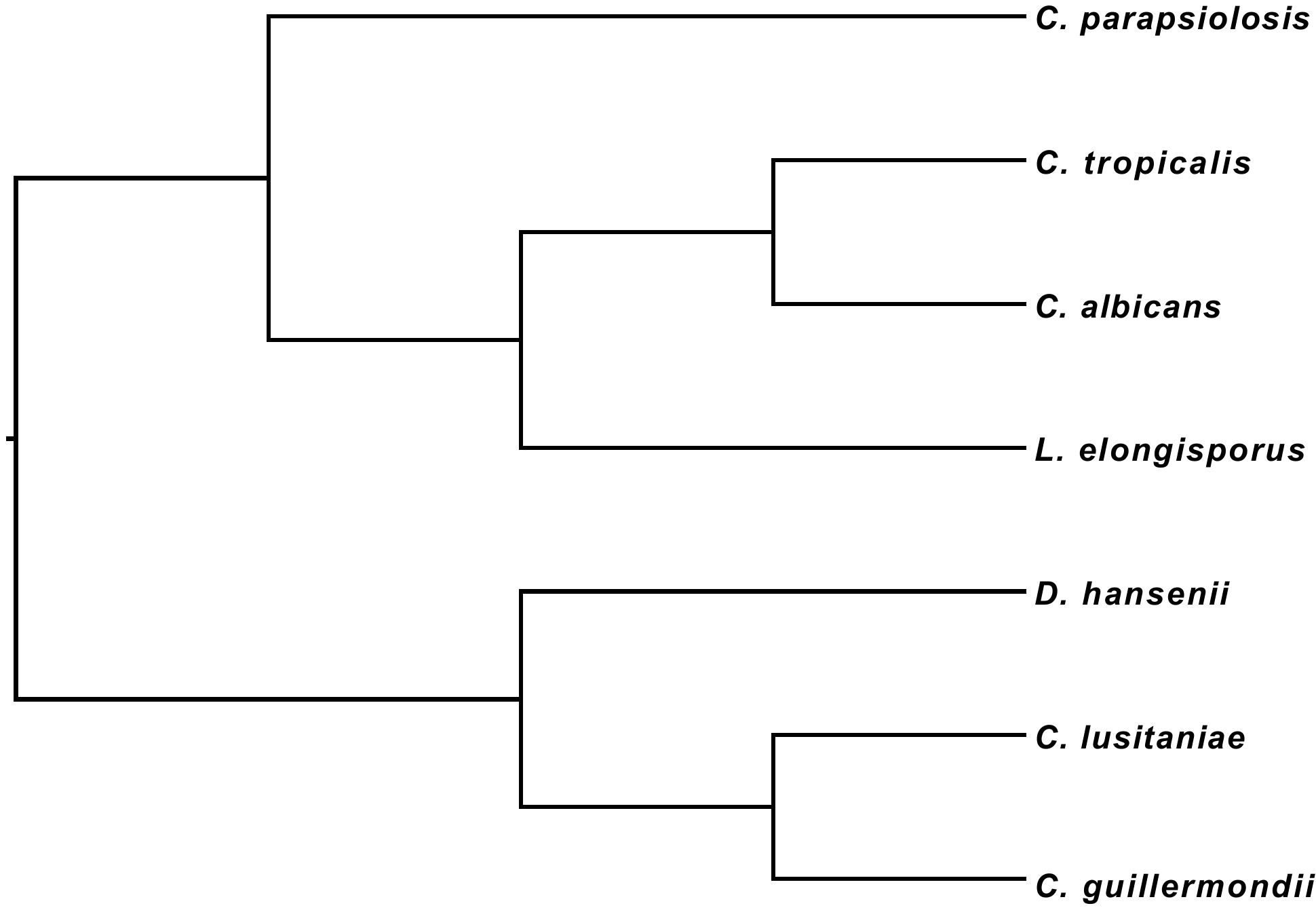

**Supporting Figure S13:** Fungal tree inferred when the final tree from phyloGAN was used for inference.

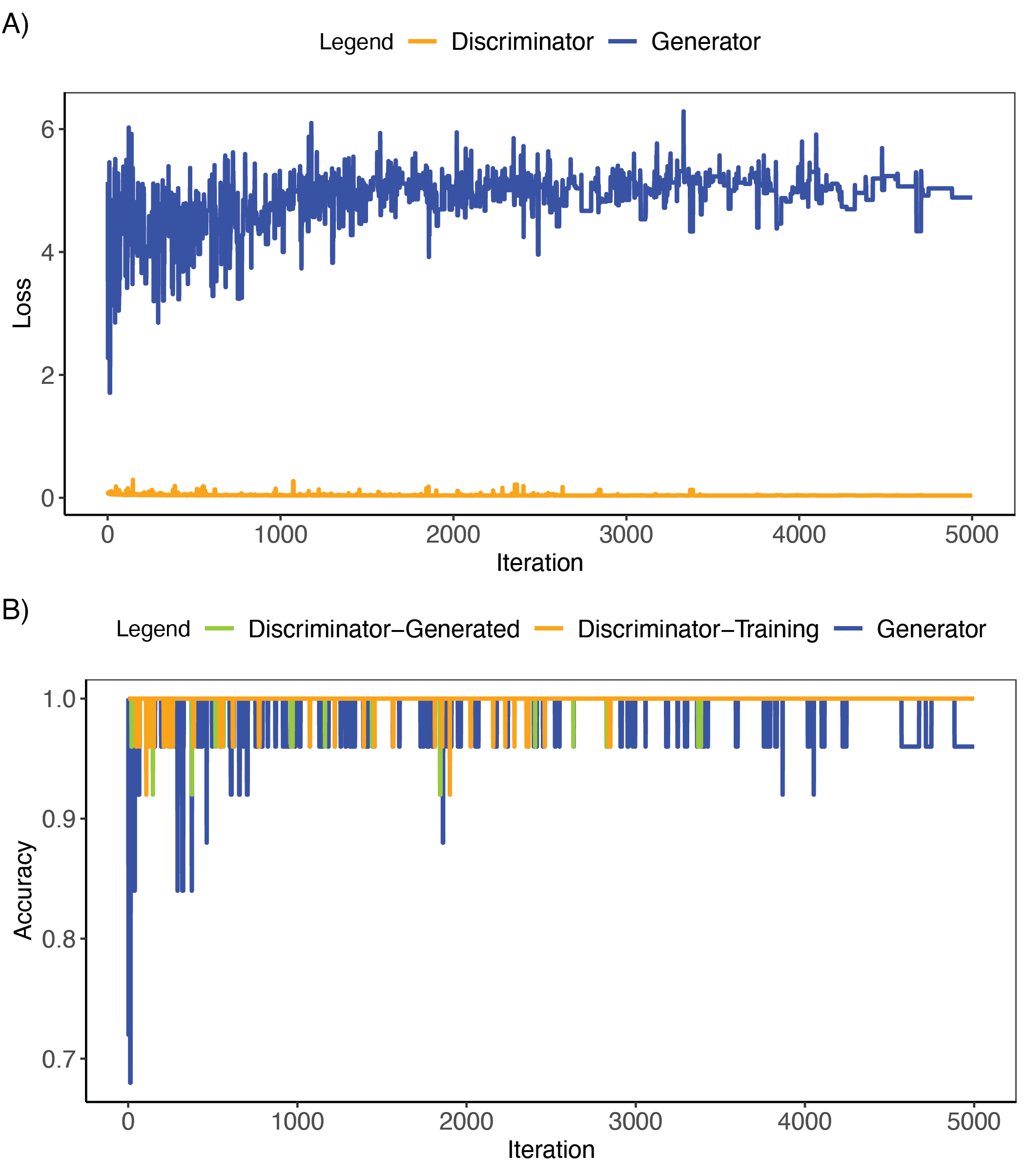

**Supporting Figure S14:** Results from analyses of yeast data. A) Generator and discriminator loss. B) Generator and discriminator accuracy. Discriminator-generated and discriminator-training indicate discriminator accuracy on generated and training data, respectively, after the discriminator was trained. Generator accuracy indicates the accuracy of the discriminator on generated data prior to training for the current iteration.

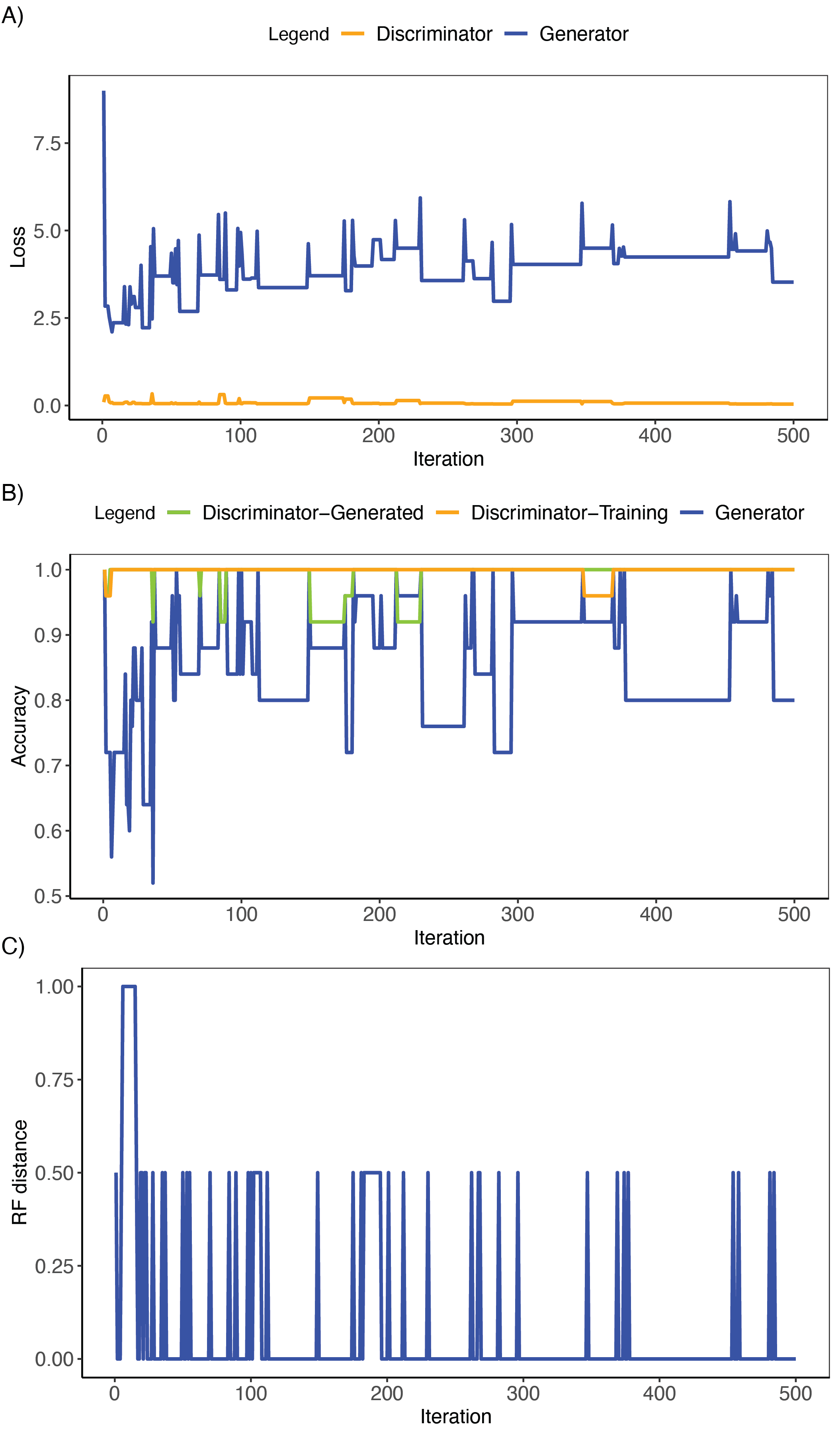

**Supporting Figure S15:** Results when branch lengths varied across the ‘observed’ dataset. A) Generator and discriminator loss. B) Generator and discriminator accuracy. Discriminator-generated and discriminator-training indicate discriminator accuracy on generated and training data, respectively, after the discriminator was trained. Generator accuracy indicates the accuracy of the discriminator on generated data prior to training for the current iteration. C) Normalized RF distances throughout the run.
